# The role of β-Nicotyrine in E-Cigarette abuse liability I: Drug Discrimination

**DOI:** 10.1101/2024.07.12.603310

**Authors:** JR Smethells, S Wilde, P Muelken, MG LeSage, AP Harris

## Abstract

**Background:** β-Nicotyrine (β-Nic) is a unique minor alkaloid constituent in electronic nicotine delivery systems (ENDS) that is derived from nicotine (Nic) degradation and can reach 25% of Nic concentrations in ENDS aerosol. β-Nic slows Nic metabolism and prolongs systemic Nic exposure, which may alter the discriminability of Nic. The present study sought to examine β-Nic has interoceptive effects itself, and if it alters the subjective effects ENDS products within a drug-discrimination paradigm.

**Methods:** The pharmacodynamics of β-Nic were examined in vitro, and a nicotine discrimination paradigm was used to determine if β-Nic (0 – 5.0 mg/kg) shares discriminative stimulus properties with Nic (0.2 mg/kg) in male (n = 13) and female (n = 14) rats after 10- & 60-min β-Nic pretreatment delays. A second group of rats was trained to discriminate β-Nic and Nornicotine (Nornic) from saline to determine if β-Nic alone has interoceptive properties and whether they are similar to Nornic.

**Results:** β-Nic had similar binding affinity and efficacy at the α4β2 nicotinic receptor subtype as Nornic, ∼50% of Nic efficacy. However, β-Nic only weakly substituted for Nic during substitution testing in female rats, but not males, whereas Nornic fully substituted for Nic. Combination testing at the 10 and 60-min pretreatment intervals showed that β-Nic dose-dependently increased the duration of nicotine’s discriminative stimulus effects, especially at the 60-min delay. Drug naïve rats could reliably discriminate Nornic, but not β-Nic, from Sal.

**Conclusion:** β-Nic increased and prolonged the interoceptive stimulus properties of Nic, suggesting it may alter to the abuse liability of ENDS through its ability to slow Nic metabolism.

## 1. Introduction

The 2022 Monitoring the Future Survey found that a greater percentage of high school seniors report past 30-day use of electronic nicotine delivery systems (ENDS) compared to cigarettes (20.7% vs 4.0%, respecitvely; Meich, et. al., 2022). This is in stark contrast to 2011 when the past 30-day use of ENDS was vastly overshadowed by combustible cigarettes (1.5% vs 15.8%, respectively; Arrazola et al., 2015). Thus, ENDS are now primarily responsible for establishing nicotine (Nic) dependence in adolescents and understanding what product charateristics of ENDS contribute to its abuse liability is critical to inform tobacco regulatory policy.

Nic is the primary constituent underlying the abuse liability of tobacco products (Benowitz, 2010; Benowitz et al., 2016), yet their addictive properties cannot be solely attributed to Nic alone. Several studies have shown that smokers who switch from conventional tobacco cigarettes (e.g., 15 mg Nic) to very-low Nic content cigarettes (VLNC; e.g., 0.5 mg Nic or %0.33 of full flavor) still maintain considerable smoking levels (15-20 VLNCs/day; Benowitz et al., 2012; Donny et al., 2007, 2015), although more recent studies show lower rates of smoking VLNCs when alternative nicotine products (e.g. ENDS) are available (Hatsukami et al., 2024). The continued use of VLNCs could be due, in part, to non-Nic constituents that are present in VLNC smoke (e.g, Nornic, acetaldehyde, β-carbolines; Won Weymarn et al., 2016). Such compounds can mimic or enhance the central nervous system (CNS) effects of Nic and/or have abuse liability themselves (Bardo et al., 1999; Benowitz et al., 2016; Caggiula et al., 2009; LeSage et al., 2018). As such, they are considered by the FDA CTP to be addiction-related harmful or potentially harmful constituents” (HPHCs; Hoffman and Evans, 2013). This designation by the FDA identifies constituents whose levels should be monitored and potentially limited to reduce the toxicity and addictiveness of tobacco products. However, this list was last updated in 2012 before ENDS rose in popularity. Given that ENDS heat a liquid rather than ignite tobacco, there is a large disparity in the chemical profile of ENDS aerosol vs combustible tobacco products. For example, β-nicotyrine (β-Nic) can be substantially more prevalent in ENDS aerosol (∼50x or 25% of Nic levels) than in tobacco smoke (Jacob et al., 1999; Martinez et al., 2015; Son et al., 2018). This is a concern because β-Nic is a potent inhibitor of the hepatic enzyme cytochrome P450 (2A6 & 2A13) and slows Nic metabolism both *in vitro* (Denton et al., 2004; Kramlinger et al., 2012; Shigenaga et al., 1989) and *in vivo* (Stålhandske & Slanina, 1982). This work prompted a <u>nicotyrine hypothesis</u> (Abramovitz et al., 2015) that it should modulate ENDS abuse liability in humans.

β-Nic is formed in ENDS through two processes: 1) dehydrogenation during ENDS aerosolization (Clayton et al., 2009) and 2) oxidation as ENDS liquid ages (Martinez et al., 2015; Wada et al., 1959). Dehydrogenation occurs during the pyrolysis (i.e., heating) of Nic-containing e-liquid solutions. Specifically, the greatest level of thermal degradation of Nic into β-Nic occurs at 200-550°C (Clayton et al., 2009; Liu et al., 1999; Woodward & Haines, 1944), which is in the temperature range (215-475°C) that an ENDS coil reaches to aerosolize e-liquids (Flora et al., 2017; Talih et al., 2019). Studies from our lab (Harris et al., 2019) and by others (Martinez et al., 2015; Son et al., 2018) have found that the aerosol produced by fresh e-liquids can contain β-Nic concentrations up to ∼13% of Nic levels, with the range varying from 1.1% to 12.9% depending on the heating temperature, ratio of propylene glycol to vegetable glycerin (the ENDS delivery vehicle), Nic concentration, vaping topography, and other factors (see Son et al., 2018). Oxidation occurs as an ENDS liquid ages, which can be accelerated by periodically opening and resealing solutions (i.e., “steeping”; a common practice among ENDS users with refillable tanks [Zhan et al., 2017]). After several weeks, concentrations of β-nic increase to 8% of Nic levels in non-aerosolized solutions and can reach 25% of Nic levels in ENDS aerosols (Martinez et al., 2015). These upper levels are ∼1000x higher than the concentration of β-Nic in combustible cigarettes which burn at ∼700-900°C (Trehy et al., 2011), a temperature far outside of the ideal range for β-Nic formation. Given its prevalence and potential to influence ENDS abuse liability, further research that directly tests the β-Nic hypothesis is needed.

Sex differences may also influence establishment of ENDS dependence. Clinical and preclinical research consistently finds that females exceed males across all phases of the Nic use disorder process, with women having more difficulty quitting smoking in adulthood (K. A. Perkins, 2001; K. A. Perkins et al., 2000; Sofuoglu et al., 1999) perhaps due to increased Nic cue reactivity (K. A. Perkins, 2001), menstrual hormone cycling (progesterone vs. estrogen) and pharmacokinetic factors (Nic metabolism is slower in women; Benowitz & Hatsukami 1998; Carroll et al., 2004; Carroll and Lynch, 2016; Carroll and Smethells, 2016; Lynch et al., 2002; Perkins et al., 1999; Pogun et al., 2017). In line with the epidemiological research, preclinical research has often found that female rats acquire quicker and maintain higher Nic self-administration (NSA) than males, especially at lower doses (Caggiula et al., 2001; Donny et al., 2000; Grebenstein et al., 2013; Li et al., 2014; Lynch, 2009; Rezvani et al., 2008; Swalve et al., 2016a; Wang et al., 2014) and that they self-administer more than males under extended access 23 hour-sessions (vs. 1-2 hr short-access ones; Flores et al., 2019). As such, the present study sought to characterize the impact of β-Nic on the abuse liability of Nic in both sexes.

The present study used a drug discrimination paradigm to determine if Nic and β-Nic produce similar interoceptive stimulus properties and if they enhance the discriminability of one another when combined (i.e., whether they feel alike and whether the combination contributes to similar CNS effects to make either drug more discriminable; see Stolerman et al., 1984). In this model, rats are reinforced with food for appropriately responding on a Nic- or Sal-associated lever following a pre-session injection of a training dose of Nic (e.g., 0.2 mg/kg) or Sal, respectively (i.e., the injection type indicates which lever produces food reinforcers). Drugs of the same class, such as Nornic and other nAChR agonists, can enhance the interoceptive effects of Nic and cause a leftward shift in the Nic dose-response curve (Desai et al., 2016; Stolerman, 1993). One of the primary neurobiological mediators of Nic discrimination is α4β2 nAChR agonism (Smith & Stolerman, 2009), and thus we also examined if β-Nic, like Nornic, has binding affinity and efficacy at this receptor subtype. If it does, it is hypothesized that like Nornic (Caine et al., 2014; Goldberg et al., 1989), β-Nic may also partially substitute for Nic during substitution testing. Regardless, the ability of β-Nic to slow nicotine metabolism is expected to prolong the half-life of Nic when they are combined and thereby shift the Nic discrimination dose-response curve to the left (Desai et al., 2016; but see Caine et al., 2014), especially under longer pretreatment intervals (10 vs 60-mins).

## 2. Method

### 2.1. Animals

Male and female (275g and 225g at delivery, respectively) Sprague-Dawley (SD; Envigo, Madison, WI) rats were individually housed with free access to chow and water in a temperature-(22° C) and humidity-controlled colony room. One week after arrival, rats were food restricted and given 16g or 18g of food per day for female and male rats, respectively. Protocols were approved by the Hennepin Healthcare Research Institute’s Institutional Animal Care and Use Committee and were in accordance with NIH guidelines set forth in the Guide for the Care and Use of Laboratory Animals (National Research Council, 2011).

### 2.2. Apparatus

Drug discrimination operant chambers (Med-Associates, St. Albans, VT) were composed of aluminum and polycarbonate walls and a stainless-steel grid floor. The chamber had two response levers on the front panel, each with a white stimulus light located directly above and a food hopper between the levers (food pellets: BioServ #F0021), and the back panel contained a house light mounted centrally at the top. Chambers were contained in sound-attenuating boxes equipped with ventilation fans that provided masking noise. A computer (OS: Windows 10^®^) running MED-PC IV^®^ (Med Associates) orchestrated experimental sessions and recorded data.

### 2.3. Drugs

(-) Nicotine base (Sigma Chemical Co., St. Louis, MO) was dissolved into saline to form a 30 mg/ml stock solution verified by gas chromatography with nitrogen phosphorous detection using our routine assay (LeSage et al., 2003). Solutions varied by no more than ± 5% (average < 1%) from the target concentration. β-Nic (obtained as a *neat oil* from Cayman Chemical, Ann Arbor, MI) was diluted to 50 mg/ml in pure EtOH. All drug doses, including Sal, were diluted in Sal and 6% v/v EtOH (w/v = 0.79g/ml; a non-behaviorally active *i.p.* dose of 39.5 mg/kg; ED_50_ of EtOH for drug discrimination is 450-560 mg/kg, see Bienkowski & Kostowski, 1998) to obtain target doses administered at 1ml/kg. The addition of 6% EtOH to Sal and all drug combinations was done to control for the highest EtOH concentration used in the final study in this series (3.0 mg/ml β-Nic *iv* self-administration, 1% higher than necessary for the present study); however, note that during substitution testing a higher 5.0 mg/kg dose was employed (i.e., 10% EtOH v/v), however, given the lack of behavioral effects at this dose the entire substitution dose-effect curve was not redetermined with this higher EtOH concentration. All drug doses are expressed as the base.

### 2.4. Procedure

#### 2.4.1. Ki determinations, receptor binding profiles and agonist/antagonist function of Nic, β*-Nic and Nornic*

Assays were conducted by the NIMH Pyschoactive Drug Screening Program (PDSP), which uses human embryonic kidney cells that stably express functional nAChR subtypes to examine the relative difference in binding affinity (assays were limited to: α2β2, α2β4, α3β2, α3β4, α4β2, α4β4 and α7) and functional agonist/antagonist efficacy (assays were limited to: α4β2 & α3β4) of Nic, β-nic and Nornic (10 mg/ml samples of each drug were submitted for analysis). The assay protocols are provided in greater detail within the PDSP assay protocol book (Roth, 2018).

##### 2.4.2.1. Nic discrimination training

Rats (n = 16 per sex; 2 females and 3 males were lost due to health issues) were initially trained (Mon-Fri) to respond in an operant chamber on two levers to earn sucrose pellets during sessions using our previously established methods (Harris et al., 2012; LeSage et al., 2012, 2009); sessions ended following 30 min or 50 reinforcers, whichever occurred first. The terminal schedule was a variable interval (VI) 15 sec schedule (for each lever, a response after an average of 15 sec produced a sucrose pellet). The active lever was alternated every other session during initial training. Once responding was sufficient on each lever (< 40 pellets per session), Nic/Sal discrimination training commenced. Specifically, rats were injected s.c. with Nic (0.2 mg/kg) or Sal (Sal; 1ml/kg) and then immediately placed in the operant chamber for a 10-min blackout prior to the start of the session. Once the training session started, as indicated by the house and cue lights illuminating, rats received sucrose reinforcers for responding on one lever if Nic was administered or the other lever if Sal was administered, with the Nic lever assignment counterbalanced across rats. Injections followed a two-session alternation between Nic and Sal (i.e., Nic, Nic, Sal, Sal, Nic, Nic, etc.). Acquisition of the Nic/Sal discrimination was assessed twice weekly (Tue & Fri) during 2-min extinction test sessions wherein no food reinforcers were delivered for Nic/Sal-appropriate responses and then the session continued normally by reinforcing drug-appropriate level responses. Once response rates and Nic discrimination during training and test sessions were stable (> 95% or > 80% injection-appropriate responding across 4 consecutive training or extinction test sessions, respectively, and no trend in overall response rate), substitution and combination testing occurred.

##### 2.4.2.2. Substitution testing and combination testing across 10- and 60-min (combination only) pretreatment intervals

Substitution and combination testing was conducted twice weekly (Tue/Fri) provided behavior was stable on intervening Nic training days (see 2.4.1.1). Following the 2-min extinction test session, rats were reinforced for responding on either lever as neither lever was drug-appropriate; the exceptions were the 0.0 and 0.2 Nic doses, which were run as normal training sessions. All ***<u>substitution testing</u>*** involved s.c. pretreatment 10 min prior to the session with Nic (0.0, 0.0125, 0.025, 0.05, 0.1, or 0.2 mg/kg), Nornic and β-Nic (0.0, 0.01, 0.025, 0.1, 0.25, 1.0, 2.5 and 5.0 mg/kg for both; drug order counterbalanced). The dose ranges of Nic and Nornic were based on prior studies showing that they bracket the range of discriminable doses (Caine et al., 2014; Stolerman et al., 2011). The lower β-Nic doses were selected to be vaping-relevant concentrations (e.g., 5 - 25% of the Nic doses used for training; Martinez et al., 2015), whereas the higher doses were based on the aforementioned Nornic drug discrimination dose range given the similarity of Nornic and β-Nic in our pharmacokinetic binding and affinity work (Fig. 1). After substitution testing, ***<u>combination testing</u>*** commenced, wherein Nic (0.0, [0.0125 10-min only], 0.025, 0.05, 0.1, & 0.2 mg/kg) discrimination was examined when combined with a low, moderate, and high dose of β-Nic (0, 0.05, 0.25, & 1.0 mg/kg, respectively) under both a 10- and 60-min drug pretreatment delay prior to the start of the session (the drug mixture was administered at 1mg/ml). The goal of the longer pretreatment interval was to assess if the ability of β-Nic to slow Nic metabolism and prolong the discriminative stimulus effects Nic. All rats were initially tested with the Nic + 0.25 mg/kg β-Nic under the 10- and 60-min pretreatment times, and then subsequent Nic + β-Nic conditions (i.e., Nic + 0.0, + 0.05 & + 1.0 mg/kg β-Nic) were counterbalanced across subjects at one pretreatment interval and then repeated at the other pretreatment interval. The Nic alone condition under the 10-min pretreatment used the data from the initial Nic dose response during substitution testing. The lower β-Nic dose (0.05 mg/kg) was chosen as it is 25% of the training Nic dose, which provided a vaping-relevant β-Nic concentrations found in ENDS vapor.

**Figure 1.**
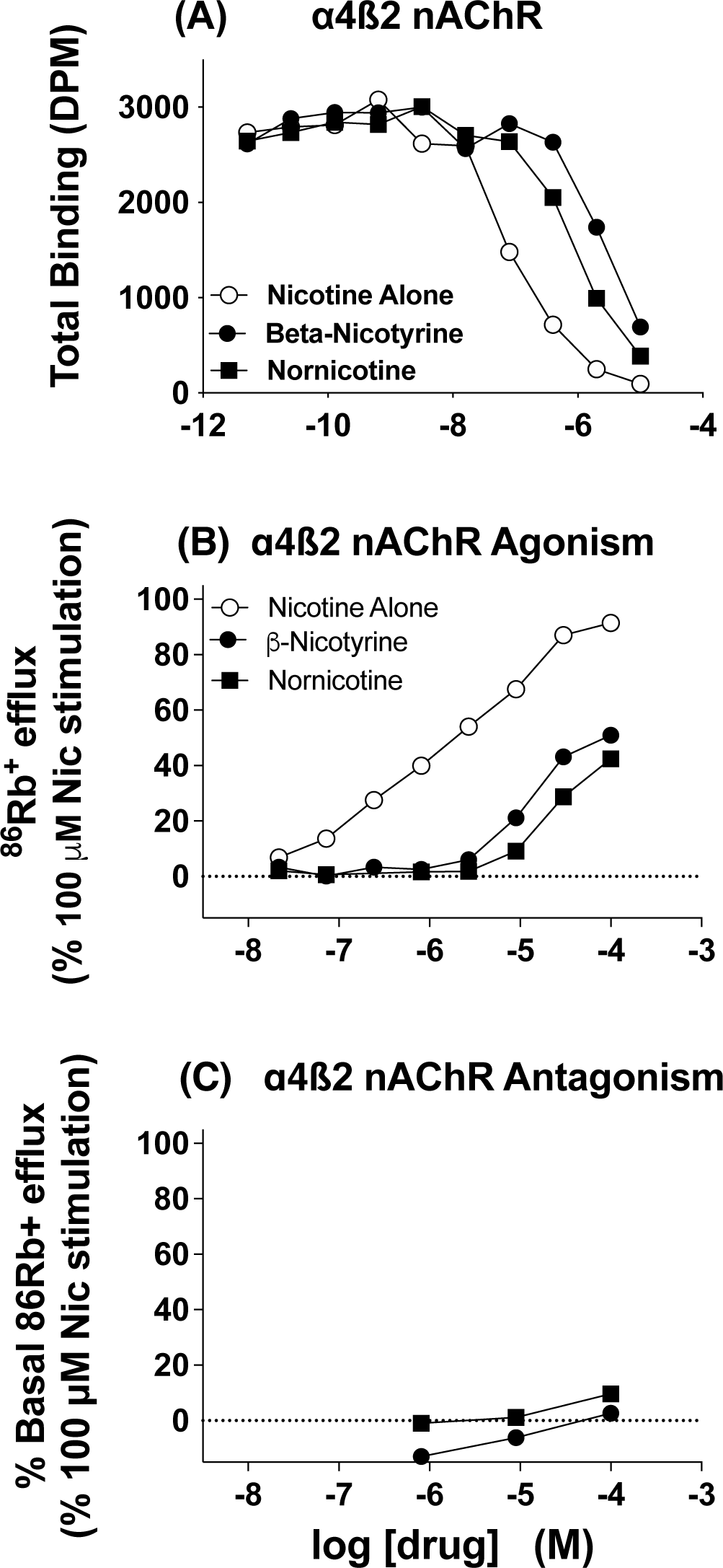
**(A)** Inhibition curve showing total nicotine radioligand binding (DPM) across a range of Nicotine, β-Nic and Nornic doses (-5 = 10μM, -6 = 1 μM, etc.). **(B)** Concentration-effect curve for stimulation of ^86^Rb^+^ efflux across a range of Nicotine, β-Nic and Nornic doses in Mols (M) as a percentage of 100 uM of nicotine. (-4 = 100μM, -5 = 10μM, etc.). **(C)** Concentration-effect curves for β-Nic and Nornic showing the percent inhibition of 100 uM nicotine on ^86^Rb^+^ efflux.

##### 2.4.2.3 β-Nic and Nornic discrimination training and testing

A separate group of rats (N = 15: 8 male, 7 female) underwent drug discrimination training using both a 2 and then a 5 mg/kg training dose of β-Nic and Nornic using the same training and testing protocol described in 2.4.1.1. Once lever press training criteria were met (reliably earning 50 pellets over 30 minutes), rats were given *sc* injections of Sal or drug 10-min prior to each 30-min session. Following two weeks of discrimination training, test sessions were conducted twice weekly (Tue/Fri) with training sessions continuing normally throughout this period. Rats experienced the following order of test sessions (min. 5 test sessions and until visually stable) that assessed discrimination between Sal (1ml/kg; *sc*) vs 2 mg/kg β-Nic (10 sessions), 2 mg/kg Nornic (10 sessions), 5 mg/kg β-Nic (10 sessions), 5 mg/kg Nornic (20 sessions), 5 mg/kg β-Nic (5 sessions).

### 2.5 Data Analysis

#### 2.5.1. Drug Discrimination

Only the data from the 2-min extinction test sessions were analyzed. The primary dependent measures were the percentage of responses on the Nic-associated lever (%NLR) and the response rate during each 2-min test session, to assess general motoric effects. The data were assessed using a mixed-model two-way ANOVA (Dose x Sex) within each drug tested and were followed by Tukey post-hoc tests to compare each drug dose to Sal, and Sidak post-hoc tests to compare drugs at each dose. Full substitution was defined as %NLR greater than or equal to 80%, while partial substitution was defined as 20%- 80% NLR. During combination testing (both at 10- and 60-min pretreatment conditions) a three-way (sex x β-Nic dose x Nic dose) ANOVA was conducted and upon a significant main effect or interaction with sex, then a separate two-way ANOVA was conducted within each sex to determine the effect of the Nic + β-Nic combination across nicotine dose. Post-hoc analyses used a Geisser-Greenhouse correction when necessary. Data analysis was conducted in Prism v.10.2.3 and R v. 4.3.2.

## 3. Results

### 3.1.1. *Ki determinations, receptor binding profiles and agonist/antagonist function of Nic,* β*-Nic and Nornic*

The inhibition constant for β-Nic was higher at the α4β2 nAChR subtype (*K*i: 103.9 nM) compared to nicotine (*K*i: 6.096 nM, Fig. 1a), but was somewhat similar to Nornic (*K*i: 69.94 nM), indicating its affinity at the α4β2 nAChR subtype is lower than nicotine and more similar to Nornic. The functional assays found that β-Nic had similar agonist properties as Nornic (Fig. 1b; i.e., they produced similar ^86^Rb^+^ efflux). Like Nornic, β-Nic did not inhibit nicotine’s agonism at the α4β2 nAChR subtype (Fig. 1c; i.e., neither produced a significant change [i.e. inhibition] in nicotine induced ^86^Rb^+^ efflux). Efficacy at the α3β4 subtype was low (9% of Nic agonism & 20% inhibition of Nic induced ^86^Rb^+^ efflux; data not shown). Lastly, binding affinity (*K*i in nM) of Nicotine, Nornic, and β-Nic was also conducted at α2β2- (Nic: 4.0; Nornic: 23.9; β-nic: 132.5), α2β4- (Nic: 6.4; Nornic: 84.5; β-nic: 173.7), α3β2- (Nic: 8.9; Nornic: 25.4; β-nic: 216.6), α3β4- (Nic: 1.95; Nornic: 2.17; β-nic: 0.6), α4β4- (Nic: 24.3; Nornic: 4.8; β- nic: 5.7) and α7-containing (Nic: 6.4; Nornic: 84.5; β-nic: 173.7) nAChR subtypes, with generally weaker affinity except at α3β4 and α4β4.

### 3.2.1. Nicotine dose-effect curve and substitution profile of β-Nic & Nornic

Figure 2 plots the %NLR and response rate for Nic, Nornic and β-Nic pretreatment 10 min prior to the test sessions. The Nic dose-effect curve showed a significant main effect of Nic dose (*_F3.31, 72.88_* = 102.3, *p* < 0.001), but no sex or sex x dose interaction, and response rates showed a sex (*_F1,25_* = 7.330, *p* < 0.05) and a sex x dose interaction (*_F5,125_* = 6.911, *p* < 0.001). Analysis of the β-Nic dose-effect substitution profile found a significant main effect of β-Nic dose (*_F7,175_* = 5.105, *p* < 0.001), with only females showing significantly higher %NLR at moderate β-Nic doses vs. Sal. There was a main effect of sex (*_F1,25_* = 8.360, *p* < 0.01) on response rate for β-Nic, but no significant within-sex effects. For Nornic (i.e., the positive control) there was a significant effect of dose (*_F7,175_* = 60.68, *p* < 0.001) and dose x sex interaction (*_F7,175_* = 2.507, *p* < 0.05) on %NLR, and a main effect of dose (*_F3.35,83.62_* = 3.854, *p* < 0.01) and sex (*_F1,25_* = 4.745, *p* < 0.05) on response rates. Post-hoc tests revealed significant differences from Sal at higher doses for both %NLR and response rate (See Figure 2).

**Figure 2.**
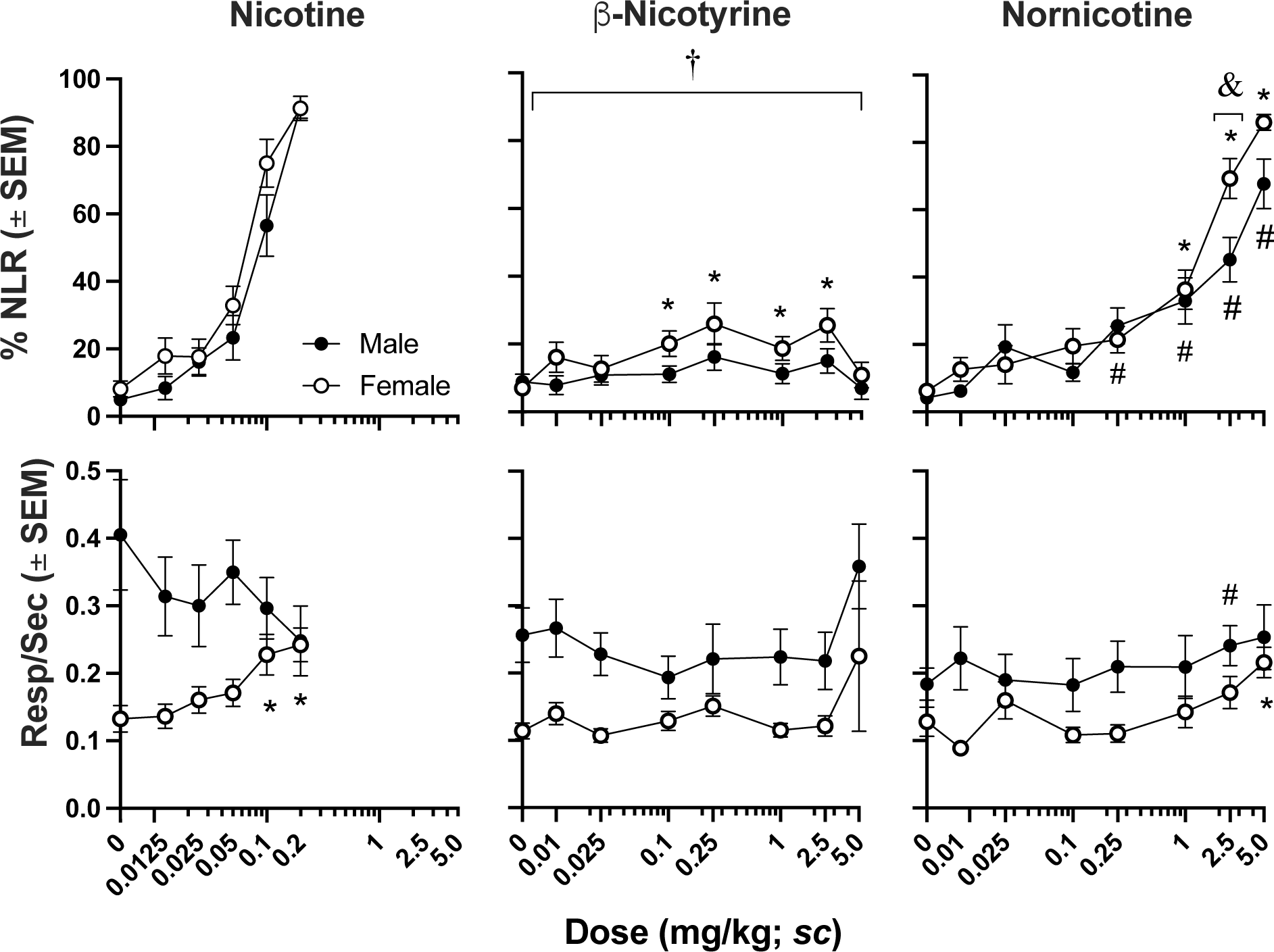
Percent nicotine lever responding and response rate (r/s) during the 2-min nicotine substation probes in male and female rats administered Nic, Nornic or β-Nic 10-min prior to the start of the session. † - indicates a main effect of sex. & - indicats interaction of sex x dose and at what dose post-hoc tests revealed differences. * and # - indicates a significant Sidak post-hoc difference from saline infemales and males, respectively.

### 3.2.2. Combination testing of Nic and β-Nic with a 10-min pretreatment interval

Figure 3 plots the %NLR for combinations of Nic + 0.05, 0.25 & 1.0 β-Nic 10 min prior to the test session. An overall analysis of the conditions found a significant effect of Sex (*_F1, 25_* = 7.98, *p* < 0.01), Nic dose (*_F5, 125_* = 418.25, *p* < 0.001), and interactions of Sex x Nic dose (*_F5, 125_* = 3.37, *p* < 0.01) and Nic dose x β-Nic dose (*_F15, 375_* = 2.04, *p* < 0.05). Analysis of the <u>Nic alone</u> condition found a main effect of Nic dose (*_F3.53, 88.19_* = 118.8, *p* < 0.001), but not sex, indicating a similar ability to discriminate Nic from saline in both sexes. At the <u>clinically relevant 0.05 mg/kg B-Nic dose</u>, there were significant main effects of Nic dose (*_F5,125_* = 203.7, *p* < 0.001), β-Nic (*_F1,25_* = 5.539, *p* < 0.05) and Sex (*_F1,25_* = 5.600, *p* < 0.05). In females, there was a main effect of β-Nic (*_F1,13_* = 5.420, *p* < 0.05), however, post-hoc tests did not show %NLR to differ at any Nic dose. At the <u>moderate 0.25 mg/kg β-Nic dose</u>, there was a main effect of Nic dose (*_F5,125_* = 201.3, *p* < 0.001), Sex (*_F1,25_* = 2.507, *p* < 0.01) and an interaction of Nic dose x Sex (*_F5,125_* = 2.332, *p* < 0.05), but no effect of β-Nic; follow-up tests found no significant main effect of β-Nic in either sex. Lastly, at <u>the highest β-Nic dose</u>, there was a main effect of Nic dose (*_F5,125_* = 262.7, *p* < 0.001), β-Nic dose (*_F0.76,18.98_* = 5.428, *p* < 0.05), Sex (*_F1,25_* = 2.507, *p* < 0.01) and interactions of Nic dose x Sex (*_F5,125_* = 2.365, *p* < 0.05) and Nic dose x β-Nic (*_F3.453,86.34_* = 3.202, *p* < 0.05). Within-sex analyses revealed a main effect of β-Nic in males (*_F1,12_* = 4.950, *p* < 0.05) and an interaction β-Nic x Nic dose in females (*_F5,65_* = 2.587, *p* < 0.05), with the combination producing significantly higher %NLR vs Nic alone at 0.1 mg/kg Nic in males and 0.05 mg/kg Nic in females (see bottom panels in Figure 3).

**Figure 3.**
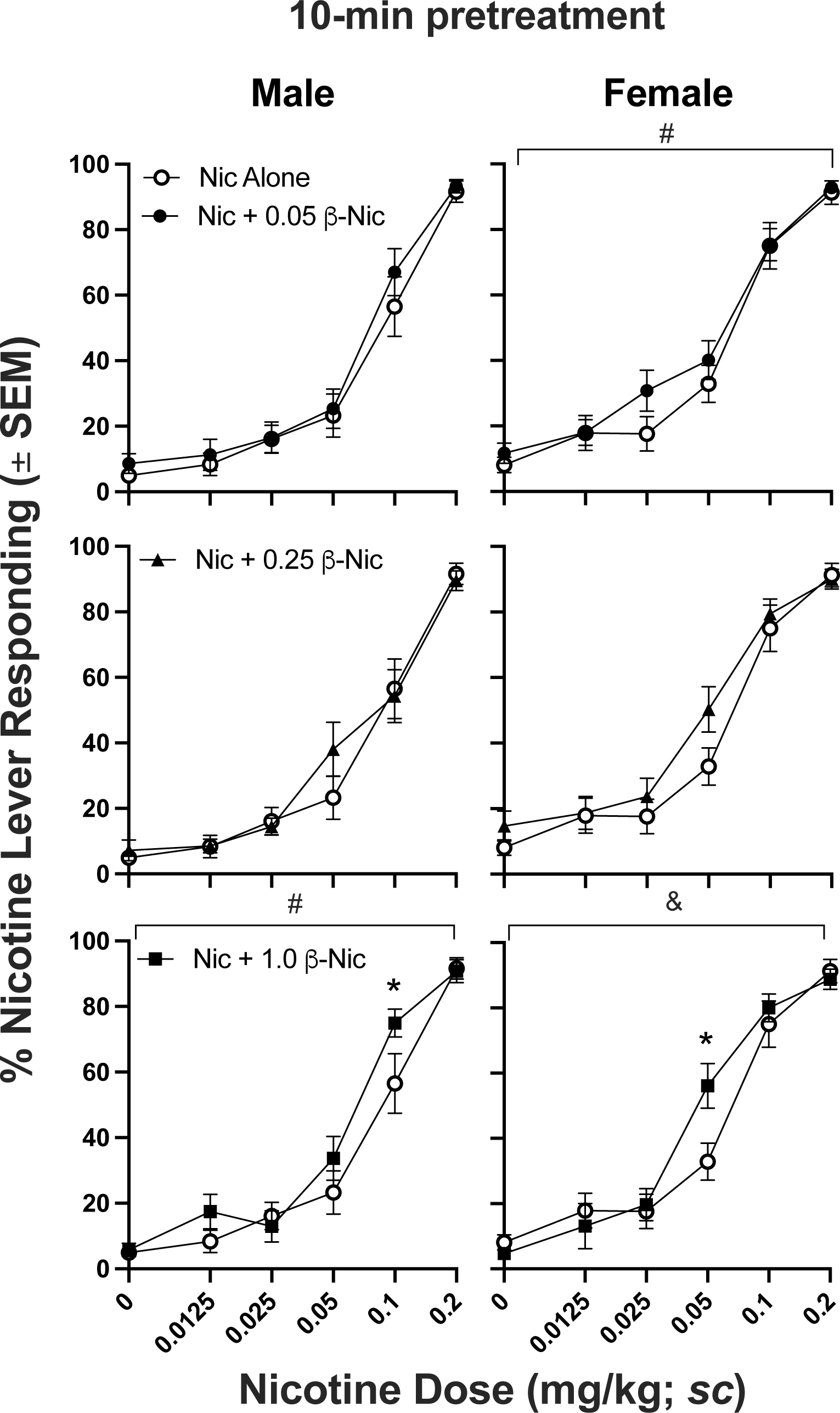
Percent nicotine lever responding during the 2-min test session as a function of Nic dose (note log axis) during Nic alone and Nic + 0.05/0.25/1.0 β-Nic under a 10-minute pretreatment schedule. Note that the Nic alone cure is the same in each panel within each sex to facilitate comparisons to combination treatments. # denotes a significant main effect of β-Nic. & denotes a significant interaction with β-Nic. * denotes a significant difference from Nic alone during post-hoc tests.

### 3.2.3. Combination testing of Nic and β-Nic with a 60-min pretreatment interval

Figure 4 displays the %NLR during the 2-min test probes that occurred 60-minutes post-injection. An overall analysis of the conditions found a significant effect of Nic dose (*_F4, 100_* = 146.5, *p* < 0.001), β-Nic dose (*_F3, 75_* = 6.59, *p* < 0.001), and interactions of Sex x Nic dose (*_F4, 100_* = 3.30, *p* < 0.05), Nic dose x β-Nic dose (*_F12, 300_* = 4.80, *p* < 0.001) and lastly a Sex x Nic Dose x β-Nic dose (*_F12, 300_* = 2.00, *p* < 0.05). In the follow-up analysis of the <u>Nic alone condition</u>, there was a significant sex difference in the ability to discriminate the interoceptive stimuli of Nic after 60 min with females showing higher %NLR than males, especially at higher doses. Analysis of the Nic Alone discrimination curves found a main effect of Nic dose (*_F2.97,74.13_* = 27.91, *p* < 0.001), sex (*_F1,25_* = 6.412, *p* < 0.05) and an interaction of Nic dose and sex (*_F4,100_* = 4.491, *p* < 0.01). Post-hoc tests revealed a significant difference at the 0.2 mg/kg dose. Analysis of the combination testing at the <u>clinically relevant dose</u>, Nic + 0.05 mg/kg β-Nic tended to show greater %NLR 60 min post injection compared to Nic alone. Analysis revealed a main effect of Nic dose (*_F4,100_* = 73.04, *p* < 0.001), β-Nic (*_F1,25_* = 7.002, *p* < 0.05), Nic dose x β-Nic (*_F4,100_* = 9.673, *p* < 0.001) and sex x Nic dose (*_F4,100_* = 4.094, *p* < 0.05); a follow-up two-way ANOVA in males and females found an interaction between Nic dose x β-Nic (*_F4,96_* = 7.002, *p* < 0.05 & *_F4,96_* = 3.597, *p* < 0.01, respectively) with both sexes showing significant differences from Nic alone at the 0.02 Nic + β-Nic combination dose in a post-hoc tests. Combination testing at a <u>moderate dose of β-Nic</u>, Nic + 0.25 mg/kg β-Nic, also found greater %NLR 60 min post injection compared to Nic alone as evidenced by a main effect of Nic dose (*_F4,100_* = 71.75, *p* < 0.001), β-Nic (*_F1,25_* = 8.614, *p* < 0.01) and an interaction of Nic dose x β-Nic (*_F4,100_* = 5.974, *p* < 0.001), β-Nic x sex (*_F1,25_* = 8.125, *p* < 0.01). A follow-up test in males found a main effect of β-Nic (*_F1,12_* = 15.05, *p* < 0.01) and an interaction of Nic dose x β-Nic (*_F4,48_* = 9.722, *p* < 0.001) with a significant difference from Nic alone at the 0.02 Nic + 0.25 β-Nic mg/kg combination dose. Analysis of the <u>highest combination</u> <u>dose of β-Nic</u> (Nic + 1.0 mg/kg β-Nic) found a main effect of Nic dose (*_F4,100_* = 83.76, *p* < 0.001), sex (*_F1,25_* = 7.354, *p* < 0.05), β-Nic (*_F0.80,20_* = 21.76, *p* < 0.001) and an interaction of Nic dose x β-Nic (*_F3.05,76.30_* = 11.32, *p* < 0.001), Nic dose x sex (*_F4,100_* = 4.374, *p* < 0.001). Follow-up tests in males found a main effect of β-Nic (*_F1,24_* = 7.813, *p* < 0.01), and an interaction of Nic x β-Nic dose (*_F4,96_* = 6.524, *p* < 0.001), whereas in females an interaction of Nic x β-Nic dose (*_F4,104_* = 4.415, *p* < 0.001) was found; post-hoc tests indicated a significant differences from Nic alone in both sexes at the 0.1 and 0.2 Nic + β-Nic combination doses.

**Figure 4.**
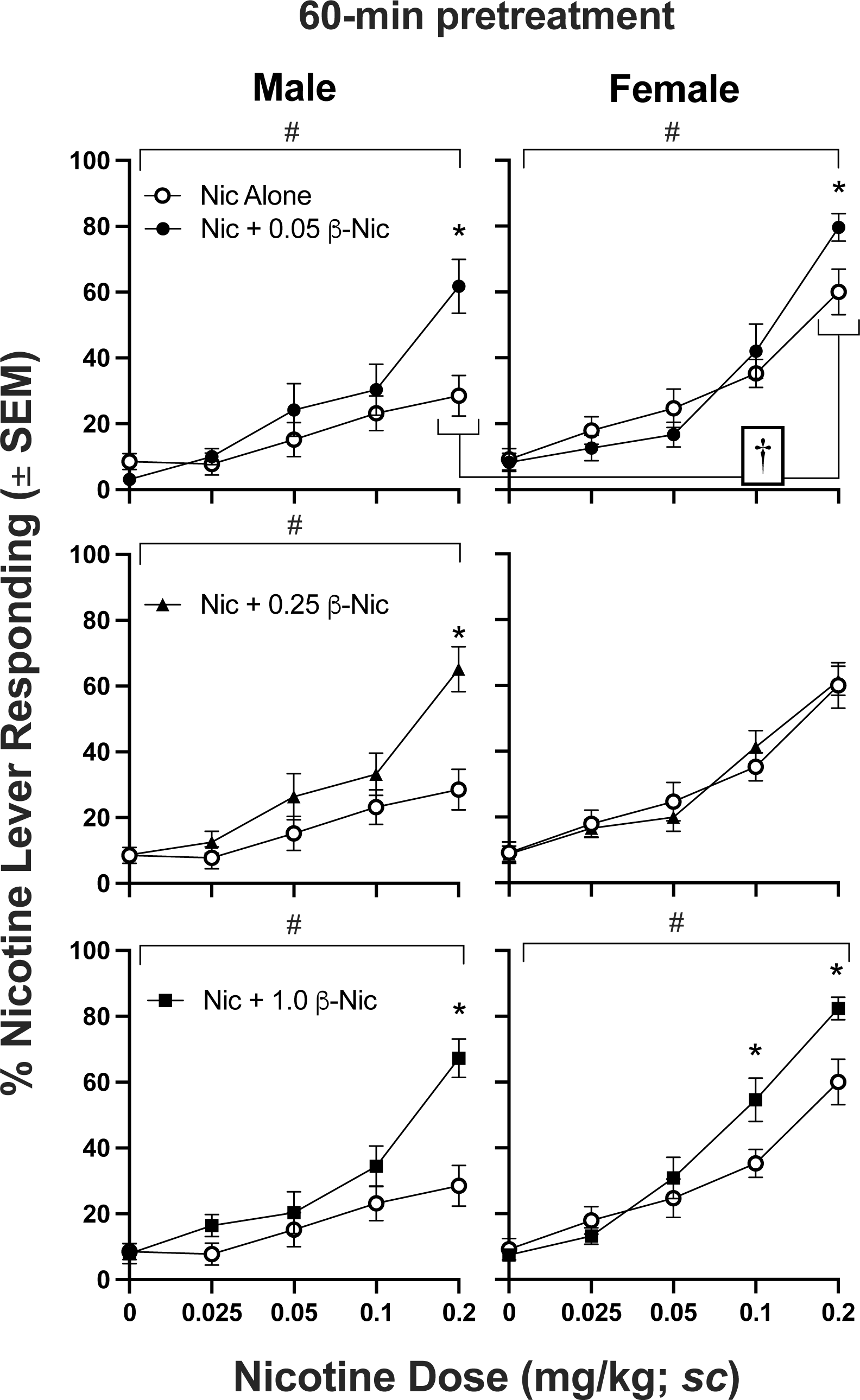
Percent nicotine lever responding during the 2-min test session as a function of Nic dose (note log axis) during Nic alone and Nic + 0.05/0.25/1.0 β-Nic under a 60-minute pretreatment schedule. # denotes a significant main effect of, or an interaction with β-Nic. * denotes a significant difference from Nic alone during post-hoc tests. † - indicates a significant difference between sexes during post-hoc tests of the Nic alone condition.

Figure 5 presents response rates during the drug discrimination sessions at both the 10- and 60-min pretreatment intervals. An overall assessment of the 10-min pretreatment condition found a main effect of Sex (*_F1, 25_* = 8.58, *p* < 0.01) and β-Nic dose (*_F3, 75_* = 3.29, *p* < 0.05), and an interaction of Sex x Nic dose (*_F4, 100_* = 12.13, *p* < 0.001) and Nic dose x β-Nic dose (*_F15, 375_* = 2.04, *p* < 0.05). To assess Sex x Nic dose interaction, response rates were assessed in both sexes. During the 10-min pretreatment condition, there were no significant differences in males, however, females showed a main effect of Nic dose (*_F2.57,33.40_* = 17.17, *p* < 0.001) on response rates across β-Nic doses (Figure 5; left column,), with significantly higher response rates at the two highest Nic concentrations within each β-Nic dose. Given there was no main effect of β-Nic dose in either sex, response rates were collapsed across β-Nic dose to examine sex differences in response rates. A main effect of Sex (*_F1,30_* = 11.45, *p* < 0.01) and Nic dose x Sex interaction (*_F5,150_* = 9.855, *p* < 0.001) were observed with significant differences between sexes at all but the two highest Nic doses. An overall assessment of the 60-min pretreatment condition found a main effect of Sex (*_F1, 25_* = 5.42, *p* < 0.05) and β-Nic dose (*_F3, 75_* = 5.67, *p* < 0.001), and an interaction of Sex x Nic dose (*_F4, 100_* = 5.87, *p* < 0.001) and Sex x β-Nic dose (*_F3, 75_* = 3.24, *p* < 0.001). To assess Sex x Nic dose interaction, response rates were assessed in both sexes. In males, there was a significant main effect of β-Nic dose (*_F2.03,24.35_* = 6.365, *p* < 0.01), but no significant differences from Nic alone were found during post-hoc tests. Females (Figure 5; right column) showed a main effect of Nic dose (*_F2.50,32.46_* = 8.002, *p* < 0.01) across β-Nic doses, with significantly higher response rates at the highest three Nic concentrations within Nic alone dose-effect.

**Figure 5.**
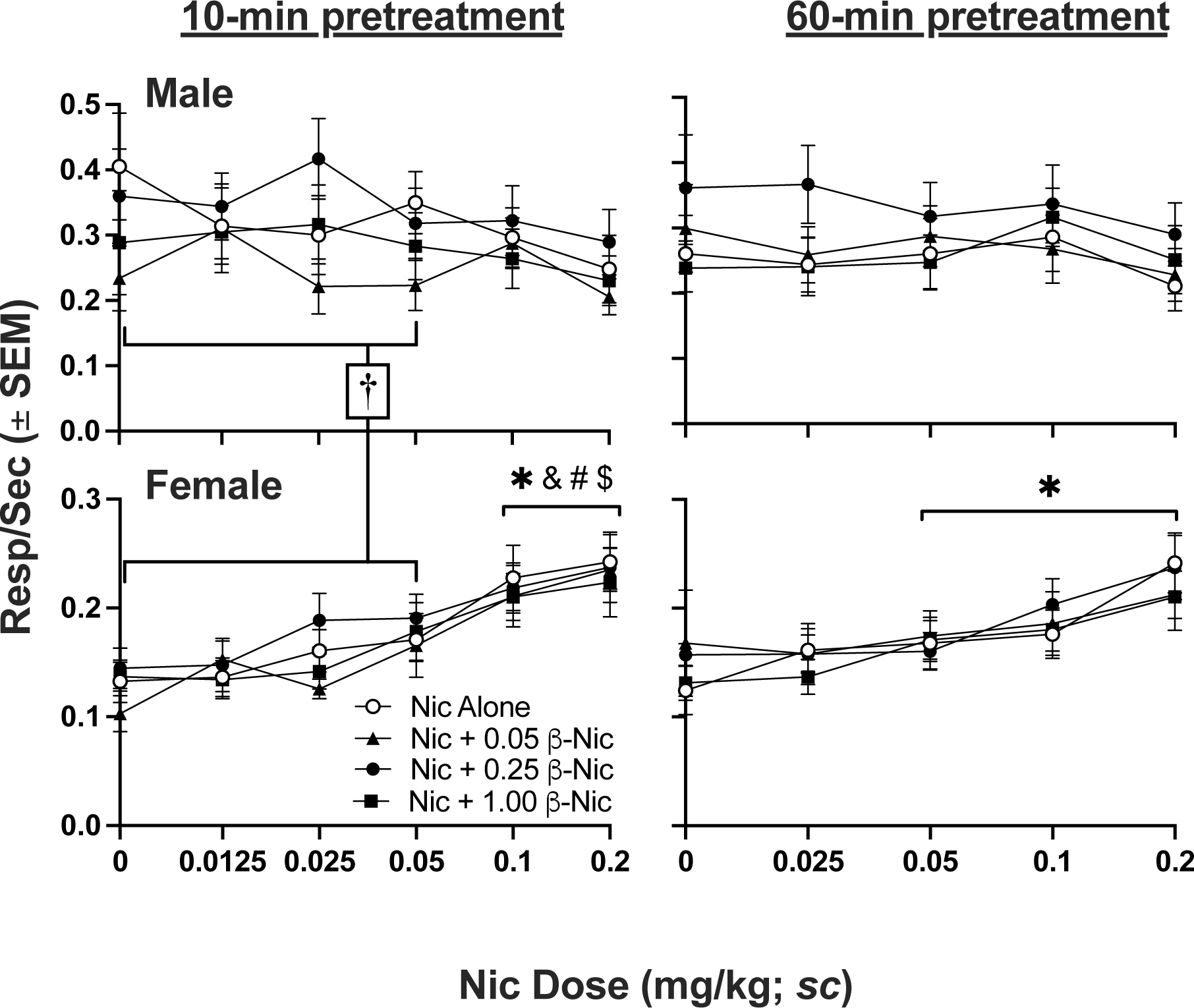
Response rates (responses/second; r/s) during the 2-min test session across β-Nic doses under the 10- and 60-min pretreatment intervals in male and female rats. *&#$ each denote a significant difference from Sal from Nic alone and Nic + 0.025/0.05/1.0 β-Nic combination doses, respectively, during post-hoc tests. † - indicates a significant difference between sexes across the bracketed Nic doses collapsing across β-Nic dose.

#### 3.3. Acquisition of β-Nic and Nornic drug discrimination

Figure 6 – left panel - shows that rats were unable to discriminate the interoceptive effects of β-Nic from Sal during discrimination testing, however, some rats were able to discriminate Nornic from Sal at both the 2 and 5 mg/kg doses (Figure 6 – right panel). Specifically, a *t*-test comparing B-Nic vs Nornic used the average of the final 3 sessions of each condition and found Nornic discrimination was only significantly higher than β-Nic at the 5 mg/kg doses (t_14_ = 7.17) after a Bonferroni correction (*p* < 0.025).

**Figure 6.**
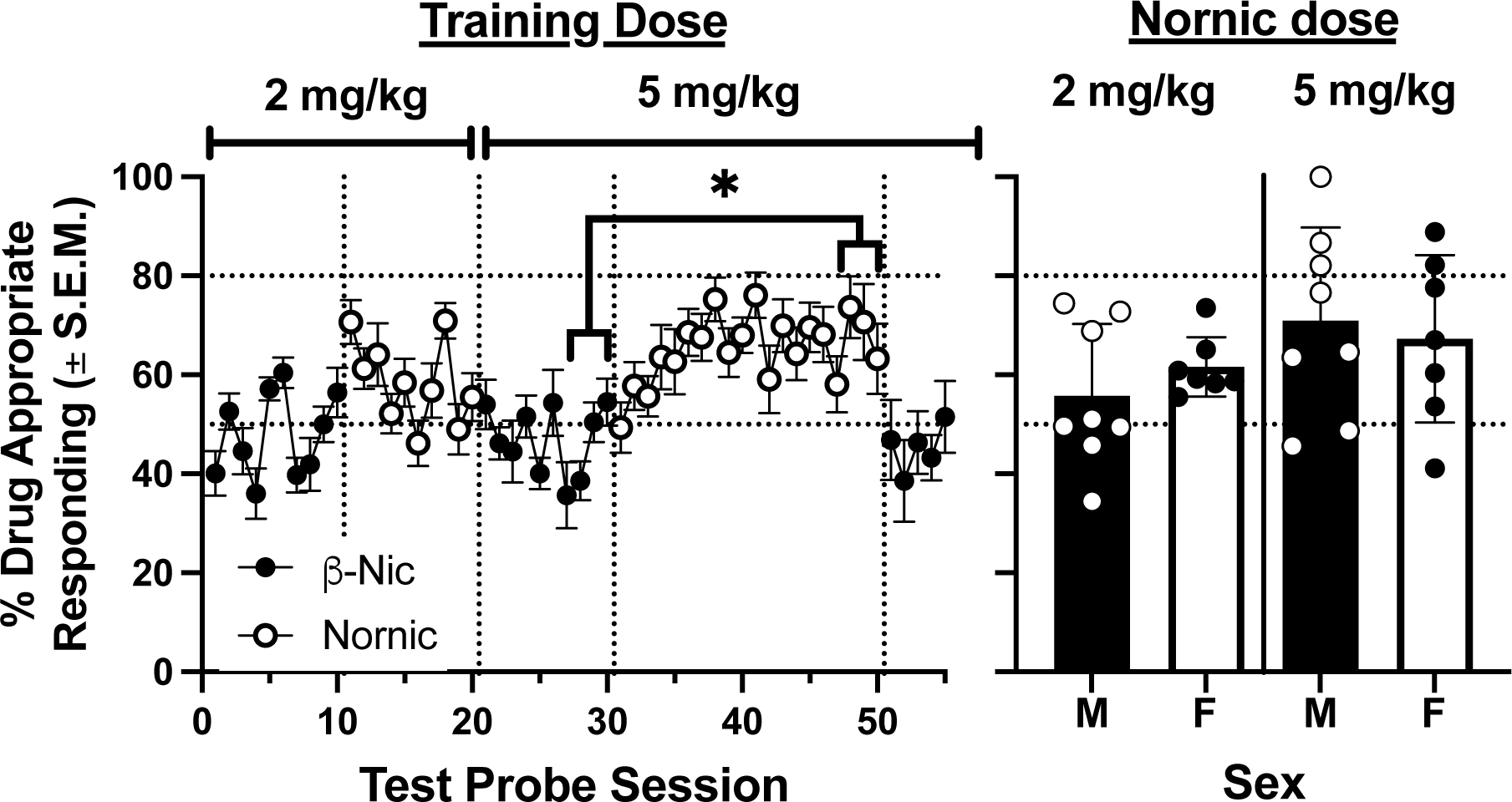
The **left panel** shows the percent drug appropriate lever responding across training sessions with 2 and 5 mg/kg training dose of β-Nic and Nornic and the **right panel** shows the individual averages of the last three sessions in each Nornic training session in male and female rats. * denotes significant difference between Nornic and β-Nic during the final 3 training sessions.

## 4. Discussion

The results of this study indicate that β-Nic prolongs the discriminative interoceptive properties of Nic in rats within a drug discrimination paradigm. The primary mechanism underlying this effect is presumed to mostly be due to the ability of β-Nic to significantly slow Nic metabolism, as evidenced by the leftward shift in the dose-effect curve during the 60-min pretreatment interval (Fig. 4). A second possible mechanism is that β-Nic has direct CNS effects, as evidenced by the PD binding/affinity at α4β2 nAChRs that was similar to Nornic, (Fig. 1), the ability for it to weakly substitute for Nic during substitution testing in females (Fig. 2), and the slightly better Nic discrimination at moderate doses when combined with β-Nic at the 10-min pretreatment interval (Fig. 3). However, the inability for β-Nic to serve as a discriminative stimulus in Nic naïve rats (Fig. 6) and the inability of it to substitute for Nic in male rats (Fig. 2), contradicts the notion that the direct CNS effects of β-Nic played a role in moderating Nic’s discriminative stimulus effects. Regardless of the mechanisms involved, the present work found that clinically relevant doses of β-Nic increase the ability of rats to discriminate Nic and may therefore contribute to the abuse liability of ENDS products.

### 4.1. Pharmacodynamics of β-Nic

The binding affinity and efficacy of β-Nic (Fig. 1) was akin to Nornic at the α4β2 nAChR. These findings are intriguing because the α4β2 subtype is the primary neurobiological mediator of nicotine’s discriminative, ICSS and reinforcing effects (Ator and Griffiths, 2003; Benowitz, 2010; Smith and Stolerman, 2009). Thus, it is surprising that β-Nic did not exhibit discriminative stimulus effects similar to Nornic and will be discussed in more detail below.

### 4.2. Substitution for Nic by β-Nic and Nornic

The present study found that Nornic almost completely substituted for Nic in both male and female rats (Figure 2), which replicates previous studies showing that Nornic has interoceptive stimulus effects similar to Nic (Caine et al., 2014; Goldberg et al., 1989). β-Nic, however, only weakly substituted for Nic in female rats, suggesting it may have weak Nic-like interoceptive properties. This finding is somewhat surprising, given that the binding and functional effects were similar to Nornic and it is presently unclear why β-Nic had no behavioral effects when administered alone.

### 4.3. Combination testing under 10- and 60-min pretreatment intervals

Combinations of Nic and β-Nic caused Nic to be more discriminable at moderate nicotine doses (Figure 3). At the 10-min pretreatment interval, this effect on Nic discrimination at moderate Nic doses was dependent on the β-Nic dose, with both sexes showing significant enhancement at the 1.0 m/kg β-Nic dose. While it is unclear whether this effect was due to the direct CNS effects of β-Nic or the ability of β-Nic to slow Nic metabolism, it does stand to reason that this effect was more likely due to the weak agonist-like CNS effects of β-Nic at α4β2 nAChRs, as a 10-min pretreatment and the session duration was likely insufficient for β-Nic’s effects on Nic metabolism to manifest. To address this issue, a 60-min pretreatment interval was employed for combination testing. Under the 60-min pretreatment interval, rats of both sexes were better able to discriminate Nic when it was co-administered with β-Nic, especially at higher β-Nic doses.

These findings imply that the interoceptive effects of Nic were prolonged by the ability of β-Nic to slow Nic metabolism by inhibiting CYP450 liver function (Kramlinger et al., 2012). During the nicotine alone condition at the 60-min pretreatment interval, female rats were better able to discriminate the training dose of Nic (0.2 mg/kg) compared to males (Figure 4 – top two panels). Response rates again showed a rate-dependent effect akin to substitution testing (Figure 5), wherein females had significantly lower baseline response rates than males that were dose-dependently increased by Nic.

### 4.4. Acquisition of Nornic and β-Nic Drug Discrimination in Nic naïve rats

The inability of β-Nic to serve as a discriminative stimulus in naive rats (Figure 6) is somewhat consistent with the Nic substitution data (Figure 2). Both male and female rats were unable to discriminate injections of 2 and 5 mg/kg of β-Nic vs Sal, whereas they were able to discriminate the higher dose of Nornic with some (%33) achieving reliable discrimination (i.e., >80%). These data call into question the reliability of the female β-Nic data showing that it can weakly substitute for Nic. This discrepancy may be due to differences in drug discrimination training history, wherein when trained with Nic female rats may be more sensitive to the interoceptive stimulus effects of β-Nic.

### 4.5. Scientific & Regulatory Implications

Teenagers are more prone to develop Nic dependence than adults (Adriani et al., 2002; Breslau and Peterson, 1996; DiFranza et al., 2002; Krishnan et al., 2022; Patton et al., 1998). Slower Nic metabolism is a risk factor for Nic dependence in naïve adolescents initiating cigarette smoking (Karp et al., 2006; for a review see Ray et al., 2009). Because slower metabolism results in prolonged systemic Nic exposure, it stands to reason that β-Nic could facilitate development of ENDS dependence in adolescents. Studies in humans are needed to examine this issue. If apparent, the FDA CTP could add β-Nic to its list of HPHCs and require manufacturers to report β-Nic levels in ENDS liquids. If necessary, a β-Nic product standard could be set to prevent increasing β-Nic levels in ENDS.

The present findings show that β-Nic may slow Nic metabolism sufficiently to increase the potency of Nic (e.g., by lowering its discrimination threshold). Prior human laboratory work by Perkins, Sofuoglu, and colleagues has examined the reinforcement and discrimination thresholds for Nic (i.e., see reviews by Perkins et al., 2022, Sofuoglu & LeSage, 2012) and the general conclusions were that greater sensitivity to the interoceptive stimulus effects of Nic is associated with greater abuse liability of Nic (e.g., Perkins et al., 2018). They contend that the discrimination threshold is a stronger metric than that of the reinforcement threshold as the former is necessary for the latter to be observed (Perkins, 2022) and that it should be the target for tobacco control (Sofuoglu & LeSage, 2012). If the present findings are applicable to humans, it suggests that β-Nic should lower the Nic discrimination threshold and thus should result in greater abuse liability of ENDS products.

## Contributions

JRS conceptualized the study and experimental design. MGL, JRS, and AHR finalized the experimental protocols. SW and PM were responsible for daily conduct of the study and data collection. JRS supervised conduct of the study and analyzed the data. JRS drafted the manuscript and MGL and AH provided editorial comments and suggestions. All authors reviewed drafts of the manuscript and approved the final version.

## Acknowledgements

We would like to thank Sam Howard and Jennifer Vigliaturo for their excellent technical assistance. Ki determinations, receptor binding profiles, agonist and antagonist functional data was generously provided by the National Institute of Mental Health’s Psychoactive Drug Screening Program, Contract # HHSN-271-2018-00023-C (NIMH PDSP). The NIMH PDSP is Directed by Bryan L. Roth at the University of North Carolina at Chapel Hill and Project Officer Jamie Driscoll at NIMH, Bethesda MD, USA. For experimental details please refer to the PDSP web site https://pdsp.unc.edu/ims/investigator/web/.

